# Genetic vitamin B6 deficiency and alcohol interaction in behavior and metabolism

**DOI:** 10.1101/2025.03.06.641947

**Authors:** Benjamin Wang, Wenqin Fu, Atsushi Ueda, Hardik Shah, Chun-Fang Wu, Wanhao Chi, Xiaoxi Zhuang

**Affiliations:** Department of Neurobiology, University of Chicago, Chicago, IL 60637; Department of Biology, College of Liberal Arts and Sciences, University of Iowa, Iowa City, IA 52242; Biological Science Division, Metabolomics Platform, Comprehensive Cancer Center, The University of Chicago, Chicago, IL 60637; The Ken & Ruth Davee Department of Neurology, Feinberg School of Medicine, Northwestern University, Chicago, IL 60611

**Keywords:** alcohol toxicity, vitamin B6, pyridoxal-5’-phosphate (PLP), pyridox(am)ine-5’-phosphate oxidase (PNPO), addiction

## Abstract

Alcohol abuse is a leading cause of preventable deaths, affecting brain function and metabolism, including GABA transmission and vitamin B6 (VB6) levels. However, the interaction between genetic VB6 deficiency and alcohol consumption remains unexplored. Here, we utilized *Drosophila* models with mutations in pyridox(am)ine-5’-phosphate oxidase (PNPO), a key enzyme in VB6 metabolism, to examine this interaction at behavioral and biochemical levels. Our findings demonstrate that PNPO deficiency reduces alcohol aversion, increases consumption, and alters locomotor behavior. Biochemically, PNPO deficiency and alcohol exposure converge on amino acid metabolism, elevating inhibitory neurotransmitters GABA and glycine. Moreover, both PNPO deficiency and alcohol exposure lead to lethality with significant interaction, which can be rescued by VB6 supplementation. These results highlight a functional interaction between genetic VB6 deficiency and alcohol, suggesting potential therapeutic strategies for alcohol-related behaviors.

## Introduction

Altered GABAergic neurotransmission and specific vitamin deficiencies are both well-documented in humans and animal models of acute and chronic alcohol abuse^1–5^. However, the potential interplay between these two effects in exacerbating alcohol toxicity and addictive behavior remains unexplored.

Ethanol (EtOH), a two-carbon alcohol, is a known allosteric positive modulator of GABA_A_ receptors, increasing the frequency and duration of GABA-gated channel opening and promoting GABA release^6–8^. The relationship between alcohol and GABAergic signaling is bidirectional. Pharmacological modulation of GABAergic signaling can directly influence alcohol intake^9–11^, and quantitative trait locus (QTL) mapping has also identified GABA-related genes associated with alcohol use^4,12–14^.

Beyond its direct effects on GABAergic signaling, chronic alcohol consumption induces deficiencies in a variety of vitamins, including vitamin B6 (VB6)^1,3^. In humans, dietary inactive VB6 (primarily pyridoxine) is converted by pyridoxal 5’-phosphate oxidase (PNPO) into pyridoxal 5’-phosphate (PLP), a critical cofactor for glutamate decarboxylase (GAD) and GABA transaminase (GABA-T)^15,16^. GAD and GABA-T are two enzymes responsible for GABA synthesis and degradation, suggesting that VB6 deficiency may contribute to alcohol-induced disruptions in GABAergic signaling.

Human genetic studies have identified severe PNPO mutations as causative for neonatal encephalopathies, and PNPO has recently been recognized as one of 16 major risk genes for adult epilepsy^17,18^. This suggests that mild PNPO mutations may contribute to epilepsy susceptibility through genetic and environmental interactions. Whether mild PNPO deficiency contributes to other disorders remains unknown. Given the complex relationship between alcohol and VB6, here we investigate the role of PNPO deficiency in alcohol use disorder (AUD).

*Drosophila melanogaster* is a well-established model for alcohol research^19–23^. Previously, we generated knock-in fly models in which the wild-type *PNPO* gene (*sugarlethal*, *sgll*) was replaced with human *PNPO* variants carrying mutations of varying enzymatic severity^24^. These models provide a unique opportunity to examine PNPO deficiency and GABAergic signaling in alcohol-related behaviors and toxicity. Using these *PNPO* knock-in flies, we identified biochemical and behavioral interactions between PNPO deficiency and alcohol exposure, providing new insights into the intersection of VB6 metabolism, GABAergic function, and alcohol-related phenotypes.

## Results

### Acute alcohol exposure reduces PLP levels in control and PNPO-deficient flies

Chronic alcohol consumption is well-documented to cause vitamin B6 (VB6) deficiency in humans and animal models^1,3^. However, the effects of acute alcohol exposure on VB6 levels remain unclear. It is also unknown whether PNPO deficiency influences this acute response. To address these questions, we measured VB6 vitamer levels, including PLP, pyridoxal (PL) and pyridoxine (PN), as well as VB6 metabolite 4-pyridoxate (PA), in flies subjected to acute alcohol (ethanol) exposure. We used the following fly strains: *w^1118^* (control)*, sgll*^95^ (a severe PNPO-deficient allele), and three human PNPO knock-in strains (*h^WT^*, *h^R116Q^*, *h^D33V^*)^24–26^. These human strains were generated by us previously, where the *Drosophila sgll* allele was replaced by the human wild-type (WT) or mutant *PNPO* cDNAs. The mutations, R116Q and D33V, result in PNPO proteins with mild to intermediate loss-of-function, respectively, based on previous biochemical and lifespan studies^17,24^.

Using mass spectrometry to analyze whole-body homogenates, we assessed VB6 levels before and after 1h ethanol vapor (40% v/v) exposure. Results revealed a significant reduction in PLP levels across all genotypes following alcohol exposure (ANOVA, treatment main effect *F_(4,40)_*=18.42, *p*=0.0004, **Fig. 1a**). A significant genotype effect was also observed (ANOVA, genotype main effect *F_(4,20)_*=6.94, p=0.001), primarily driven by *sgll*^95^, which exhibited significantly lower PLP levels compared to its genetic controls *w^1118^* (Tukey’s post hoc, *p*=0.03, **Supplementary Tables 1-2**). In contrast, PLP levels in *h^R116Q^* and *h^D33V^* were comparable to those in their genetic controls *h^WT^*, which is in line with our previous findings indicating that R116Q and D33V are milder mutations compared to the severe missense mutation in *sgll*^95^ ^24,25^.

**Fig. 1:**
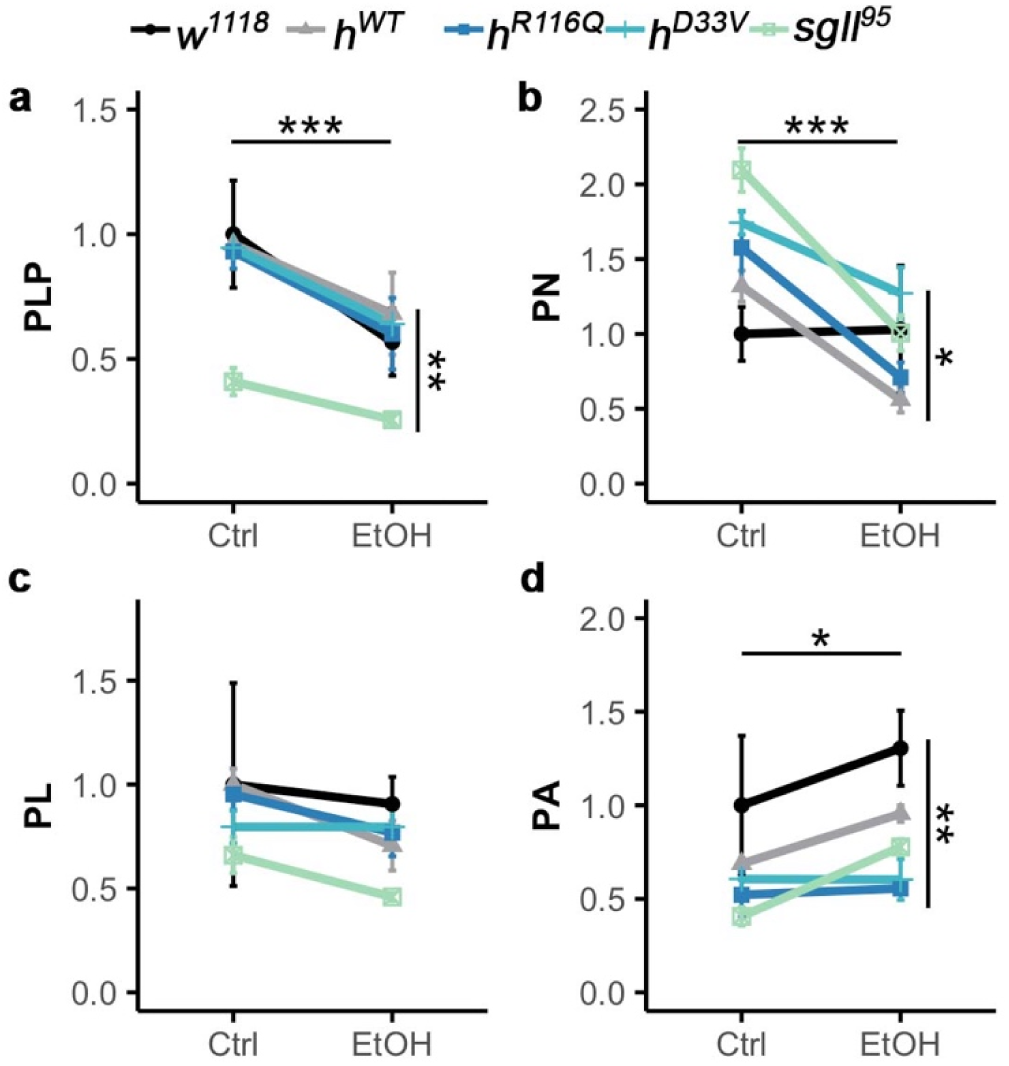
Vitamin B6 levels in control and PNPO mutant flies before and after acute alcohol exposure. Mass spectrometry quantification of tissue content of **A:** PLP (pyridoxal 5’-phosphate), **B:** PN (pyridoxine), **C:** PL (pyridoxal), and **D:** PA (4-pyridoxate, vitamin B6 metabolite) in control diet (Ctrl) or alcohol exposure (EtOH) conditions (40% v/v for 1h). *n*=3 groups per genotype per condition. Data is presented as mean ± SEM. Two-way ANOVA with Tukey’s post hoc. The horizontal and vertical bars indicate the treatment and genotype effect, respectively. *p<0.05, **p<0.01, ***p<0.001. Significance for genotype comparisons within treatment groups are included in the text.

PN levels followed a trend similar to PLP (ANOVA, treatment main effect *F_(4,20)_*=26.7, *p*=4.66e-5; genotype main effect *F_(4,20)_*=4.26, *p*=0.01, **Fig. 1b**). PL levels showed no significant changes with treatment or genotype (**Fig. 1c**). PA levels, however, increased overall following alcohol exposure (ANOVA, treatment main effect *F_(4,20)_*=4.49, *p*=0.047; genotype main effect *F_(4,20)_*=6.08, *p*=0.002, **Fig. 1d**). Notably, we found no evidence of a genotype-by-treatment interaction for these VB6 vitamers, suggesting that the acute effect of alcohol on VB6 levels is likely independent of PNPO function.

Taken together, these findings demonstrate that acute alcohol exposure accelerates VB6 catabolism, leading to a depletion of PLP levels.

### Chronic alcohol exposure impairs survival, which can be improved by PLP supplementation

Our previous studies demonstrate that severe PLP deficiency caused by PNPO mutations can result in death in flies^24–26^. Since both PNPO mutations and alcohol exposure can lead to PLP deficiency (**Fig. 1a**), we investigated their potential additive effects on survival. Additionally, we included PLP supplementation to explore its rescue effect on survival of flies from the various genotypes. Flies were introduced into vials and given *ad libitum* access to a VB6-depleted diet either lacking or containing 16% ethanol, without or with PLP supplementation.

In the condition of no alcohol and no PLP supplementation, *sgll*^95^ flies exhibited the lowest survival rate within 21 days, followed by *h^R116Q^* and *h^D33V^* flies (**Fig. 2**, the top left panel in **Fig. S1a**, **Supplementary Tables 1-2**). No death was observed in *w^1118^* and *h^WT^* flies (**Fig. 2, Fig. S1b**). Thus, survival on the VB6-depleted diet is largely correlated with the residual PNPO enzymatic activity.

**Fig. 2:**
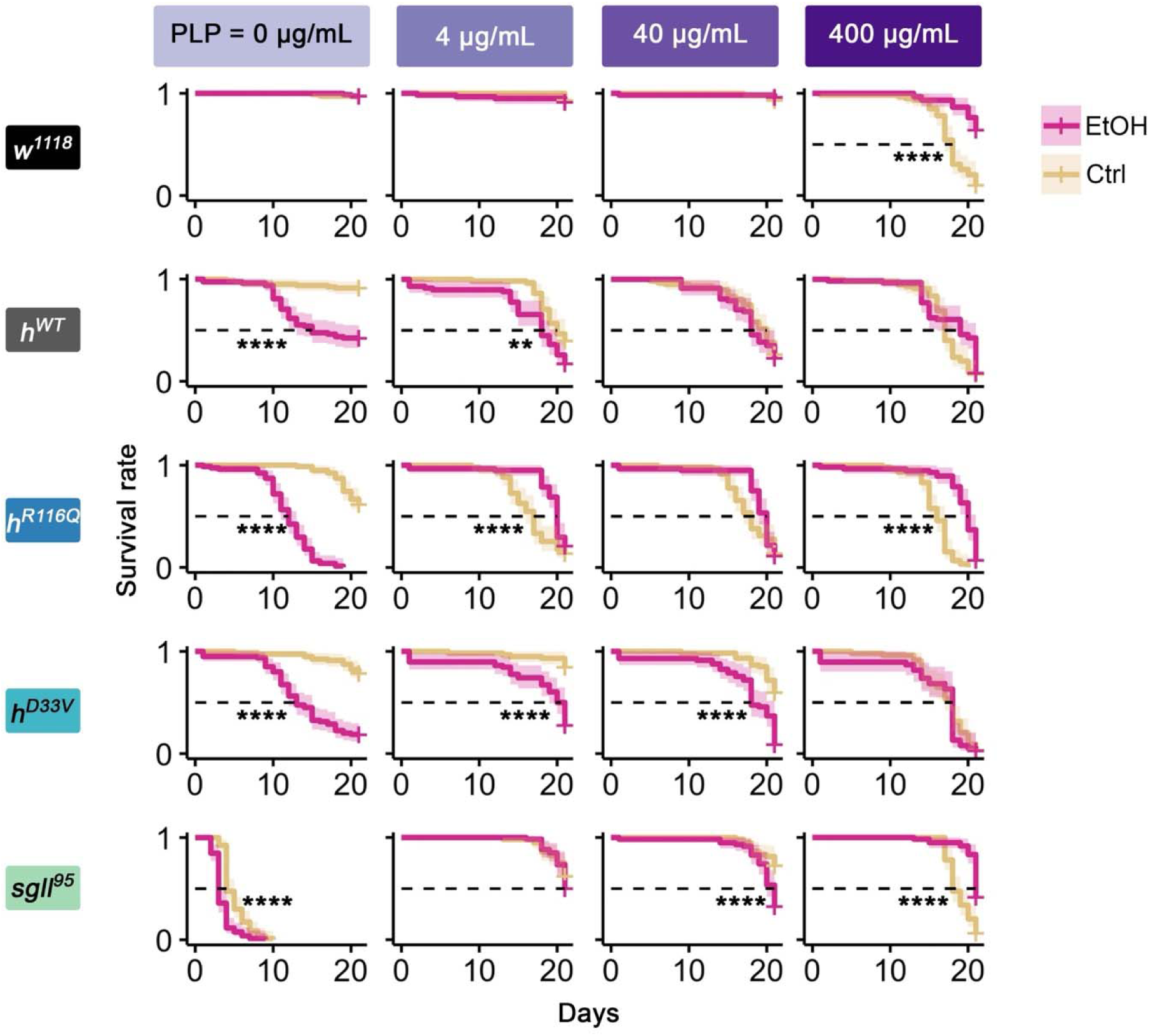
Effects of chronic alcohol exposure and PLP supplementation on survival of flies from each genotype. Survival of flies from each genotype on control diet (Ctrl) or 16% alcohol (EtOH) for 21 days in combination with PLP supplementation. *n*=38-80 flies per genotype per diet treatment. Log-rank test with Benjamini-Hochberg corrections. **p<0.01, ****p<0.0001. See Supplementary Tables 1-2 for exact n in each condition and exact p values for comparisons.

Previously, we showed that low concentrations of PLP (<400 ng/mL) rescued survival of *sgll*^95^ flies on the VB6-depleted diet in a dose-dependent manner^26^. However, excess pyridoxine, the precursor of PLP, has been associated with aldehyde toxicity and neuropathy in humans^27,28^. To test the rescue and toxicity effects of PLP on flies with varying levels of PNPO deficiency, we supplemented their diets with high concentrations of PLP (4, 40, and 400 μg/mL). Survival data indicated that PLP at 400 μg/mL induced significant toxicity in all genotypes, including *w^1118^* and *h^WT^*, except *sgll*^95^ (**Fig. 2**, the left column in **Fig. S2a, Fig. S2b**), demonstrating that high levels of PLP supplementation are indeed toxic unless that the organism already has severe deficiency in endogenous PLP.

While *h^R116Q^* and *h^D33V^* flies exhibited consistent responses to high PLP levels, their responses to low or intermediate PLP supplementation (4 and 40 μg/mL) varied depending on their degree of PNPO deficiency. For example, *h^D33V^* flies showed increased lethality only to PLP 40 μg/mL while *h^R116Q^* flies exhibited increased lethality at PLP concentrations of both 4 and 40 μg/mL (**Fig. 2**, **Fig. S2b**). Thus, flies with more severe PNPO deficiency were more likely to tolerate PLP toxicity.

Since alcohol exposure depletes PLP (**Fig. 1a**), we expected to see lethality or increased lethality from all genotypes upon alcohol treatment. Indeed, we observed lethality in *h^WT^* flies, which otherwise survived well, and increased lethality in *h^R116Q^*, *h^D33V^*, and *sgll*^95^ flies, which already exhibited lethality on VB6-depleted diet (the first column in **Fig. 2)**. Notably, this chronic alcohol exposure condition did not affect the *w^1118^* flies. The differential sensitivity of *w^1118^* and *h^WT^* flies to alcohol exposure suggested that the wild-type human *PNPO* cDNA in *h^WT^* flies did not completely replace the function of the *Drosophila sgll* allele (see Discussion).

We further hypothesized that PLP supplementation could mitigate alcohol toxicity. Although survival dynamics differed across genotypes due to their varying responses to the toxicity effect of alcohol and high doses of PLP, PLP supplementation generally enhanced the tolerance of flies to alcohol exposure (**Fig. 2**).

Taken together, these results suggest that an optimal level of PLP is crucial for survival. Both excessively high PLP concentrations from supplementation and excessively low PLP levels due to PNPO mutations or alcohol exposure are detrimental for survival. Survival as a function of PLP levels thus likely follows a bell-shaped curve. Both PNPO deficiency and alcohol exposure appear to shift this curve rightward, increasing the requirement for PLP supplementation for survival and enhancing tolerance to PLP overdose.

### Mutations in the *PNPO* gene reduce aversion to high alcohol concentration

In *Drosophila* and other animal models, sensitivity to nutritional needs and food consumption is often regulated by homeostatic mechanisms^2^^9,30^. Deviations from these mechanisms may lead to maladaptive behaviors, such as excessive intake of addictive substances like alcohol, irrespective of nutritional needs or homeostatic regulation^31,32^. This phenomenon could partially explain the increased lethality observed in mutant flies on a chronic alcohol diet (**Fig. 2**). On the other hand, adult *Drosophila* generally avoid food with >5% ethanol^33^. To investigate whether PNPO deficiency affects alcohol aversion to 16% ethanol and therefore consumption, we examined alcohol preference in mutant and control flies.

A proboscis extension reflex (PER) assay^34^ was used to assess taste response to alcohol. In this assay, an alcohol (16% ethanol+4% sucrose) or isocaloric maltodextrin 4% sucrose solution droplet was brought into contact with each fly’s labellum (**Fig. 3a**). Only flies exhibiting proboscis extension to maltodextrin (more than 95% of flies in each strain, **Fig. 3b**) were further tested for their response to alcohol. Compared to *w^1118^* or *h^WT^* flies, which exhibited low alcohol response frequencies (< 20%), mutant flies showed significantly higher responses: 40% in *h^R116Q^*, 60% in *h^D33V^*, and 80% in *sgll*^95^ flies (ANOVA, *F_(4,10)_*=38.38, *p*=4.93e-06. **Fig. 3b**, **Supplementary Tables 1-2**), suggesting that mutant flies had decreased aversion to high alcohol concentration.

**Fig. 3:**
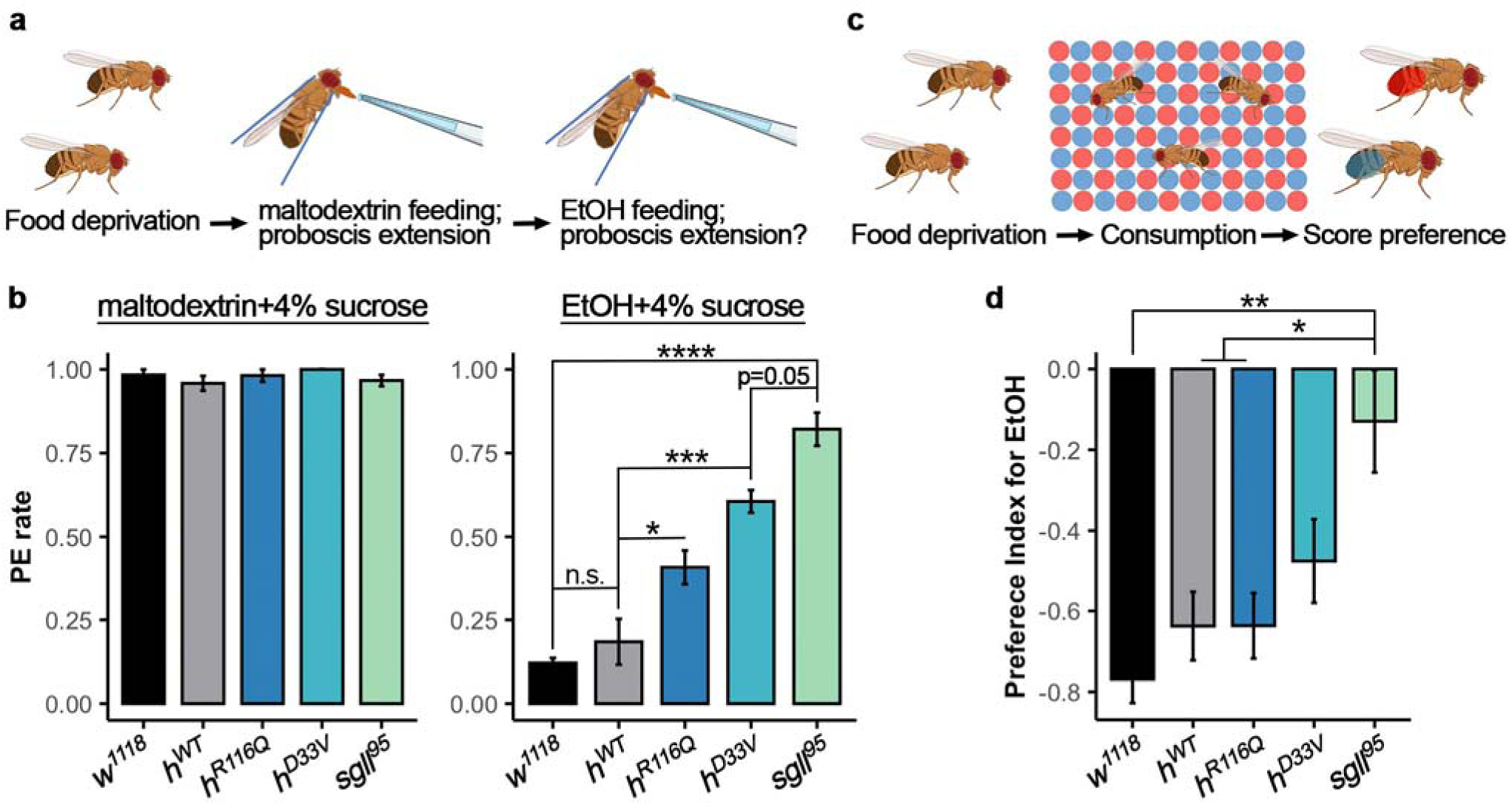
PNPO mutations decrease aversion to high alcohol concentration. **a** Schematic for proboscis extension reflex assay. **b** Proboscis extension (PE) rate for maltodextrin+4% sucrose (left) and alcohol +4% sucrose (right). *n*=3 trials per genotype. **c** Schematic for the binary food preference assay. **d** Preference Index for EtOH over isocaloric maltodextrin (see Methods for calculation). *n*=17 assays per genotype for both dye (see Supple. Fig. 3 for Preference Index for alcohol paired with blue dye or red dye). Data is presented as mean ± SEM. One-way ANOVA with Tukey’s post hoc. n.s.:p>0.05, *p<0.05, **p<0.01, ***p<0.001, ****p<0.0001. See Supplementary Table 2 for exact p values.

To confirm the decreased aversion response in mutant flies, an additional binary food choice assay was conducted^35^. Flies were given *ad libitum* access to a 96-well plate containing alternating wells of 16% ethanol + 4% sucrose or isocaloric maltodextrin + 4% sucrose in agar for 1h (**Fig. 3c**). The alcohol and maltodextrin foods were dyed blue or red, enabling determination of food consumption based on the coloration of the fly abdomens. To account for any natural preference for a specific color or taste^36–38^, the assay was repeated with alcohol-dyed either blue or red. Although the genotype effect is significant for both dyed conditions (blue: ANOVA, *F(_4,35)_*=17.7, *p*=4.99e-08; red: ANOVA, *F_(4,40)_*=25.2, *p*=1.78e-10), we found flies generally avoided food with red dye^38^ (**Fig. S3**, **Supplementary Tables 1-2**). Data averaged from both dyes showed a significant genotype effect (ANOVA, *F_(4,80)_*=4.95, *p*=0.001. **Fig. 3c**, **Supplementary Tables 1-2**), primarily driven by *sgll*^95^ flies (Tukey’s post hoc*, p*=0.001, 0.015, and 0.011, compared to *w^1118^*, *h^WT^*, and *h^R116Q^*, respectively). Together, these findings suggest that PNPO mutations decrease aversion to high alcohol concentration, which may contribute to (at least partially) the reduced survival of mutant flies under chronic alcohol exposure.

### Mutations in the *PNPO* gene affect alcohol sensitivity but not tolerance development

We next investigated alcohol sensitivity and the development of tolerance to alcohol using a repeated exposure paradigm^39^ (**Fig. 4a**). Flies from each genotype were exposed twice to 40% ethanol vapor, with exposures spaced 16 hours apart. The initial exposure lasted for 1h and the second exposure lasted for 1.5h. The time required to reach 50% sedation in each group (ST50) was used as a measure of alcohol sensitivity. Upon initial exposure, there is a significant genotype effect (ANOVA, *F_(4,33)_*=15.8, *p*=2.55e-07, **Fig. 4b-c**, **Supplementary Tables 1-2**). Specifically, *w^1118^* flies exhibited the greatest sensitivity to alcohol sedation, as indicated by a significantly shortened sedation time compared to all other genotypes (Tukey’s post hoc, *p*=1.5e-06, 0.0007, 7e-07, and 0.008, compared to *h^WT^*, *h^R116Q^*, *h^D33V^*, and *sgll*^95^, respectively. **Fig. 4b-c**). The *h^WT^* and *h^D33V^* flies were least sensitive to alcohol sedation, while *h^R116Q^* and *sgll*^95^ exhibited moderate sensitivity, although statistical analyses indicated that there was no difference among the three human mutant strains (**Fig. 4c, Supplementary Table 2**).

**Fig. 4:**
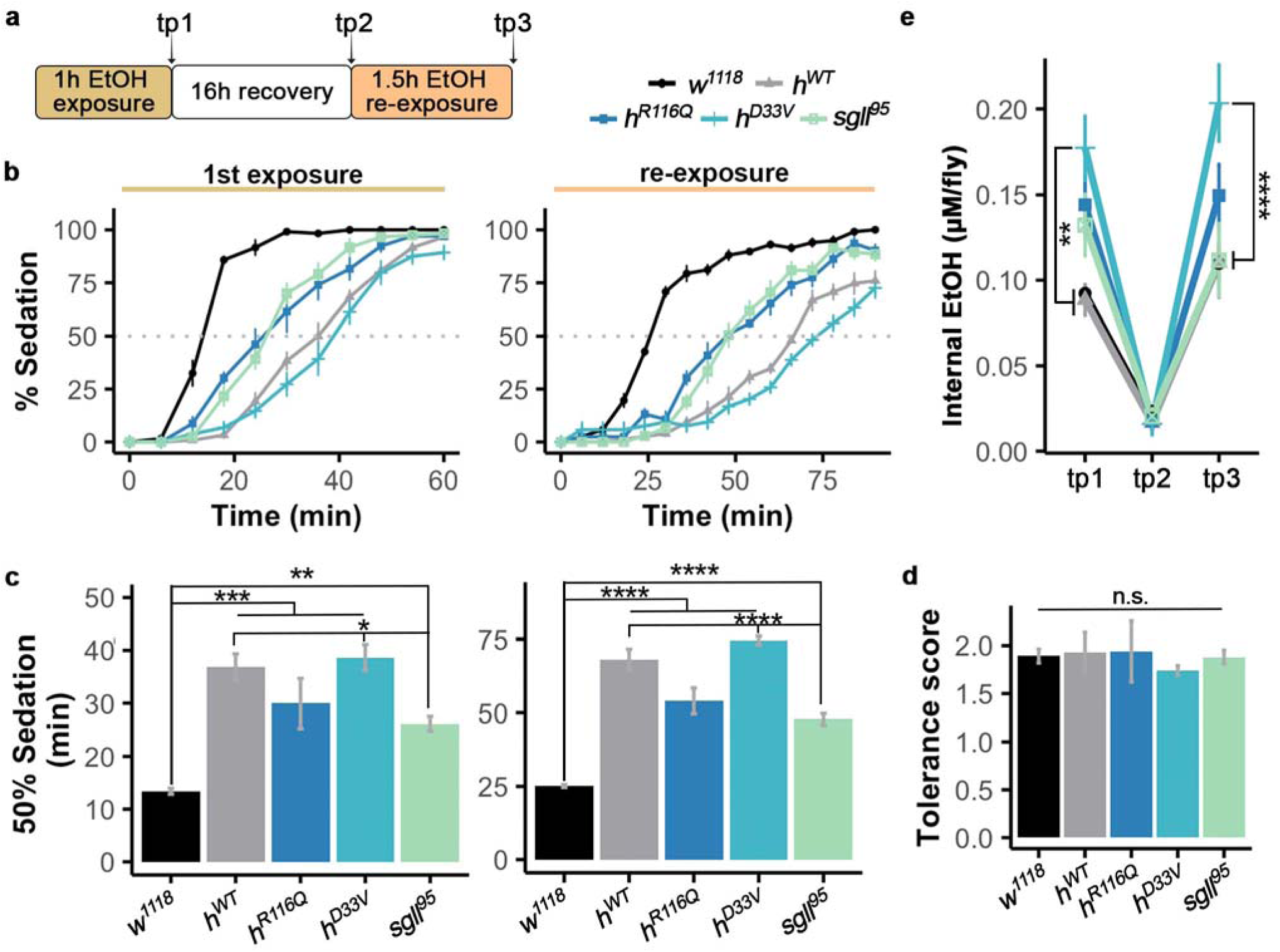
PNPO mutations affect alcohol sensitivity but not tolerance development. **a** Schematic for sensitivity and tolerance assay, tp = timepoint at which flies were collected. **b** Percentage of sedated flies upon the initial alcohol exposure (left) and the alcohol re-exposure after 16h recovery (right). *n*=8 trials. **c** Quantification of time to 50% sedation (ST50) of each genotype upon the initial alcohol exposure (left) and the re-exposure after 16h recovery (right). **d** Development of tolerance of each genotype upon re-exposure to alcohol, calculated by 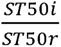(i=initial exposure, r=re-exposure). **e** Whole-body alcohol levels in flies after alcohol exposure at 3 timepoints. *n*=3 trials. Data is presented as mean ± SEM. Two-way ANOVA with Tukey’s post hoc. n.s:p>0.05, *p<0.05, **p<0.01, ***p<0.001, ****p<0.0001. See Supplementary Table 2 for exact p values.

During the second exposure, ST50s were increased for all genotypes (**Fig. 4b-c)**, indicating the development of tolerance in them. The tolerance development was comparable among all genotypes (**Fig. 4d, Supplementary Table 2**). Together, these results suggest that while PNPO mutations likely decrease sensitivity to alcohol, they do not seem to impair the ability of flies to develop tolerance to alcohol.

### Mutations in the *PNPO* gene affect alcohol clearance

We further investigated whether PNPO mutations affect alcohol clearance. Following the tolerance study, we pooled flies from each genotype and experimental timepoint to measure whole-body alcohol levels (**Fig. 4a**). Alcohol levels were assessed using a fluorescence-based detection method at three timepoints: immediately after 1h of alcohol exposure, after a 16h recovery period, and after 1.5h alcohol re-exposure following the recovery. After the initial 1h exposure to 40% ethanol vapor, *h^R116Q^*, *h^D33V^*, and *sgll*^95^ tended to have higher body alcohol levels than *h^WT^* and *w^1118^*, which are comparable (Tukey’s post hoc, *p*=0.005 for *h^D33V^*, compared to *h^WT^*, tp1 in **Fig. 4e**, **Supplementary Table 2**). The increased body alcohol level in *h^D33V^* flies was likely attributed to compromised alcohol catabolism, although we could not exclude the possibility of increased alcohol absorption in them.

Following the 16h recovery period, alcohol levels decreased significantly in all genotypes and no difference were observed among them (tp2 in **Fig. 4e**, **Supplementary Table 2**). After the 1.5h re-exposure to alcohol following the recovery period, the *h^D33V^* flies again exhibited a significant higher body alcohol level than *h^WT^* (Tukey’s post hoc, *p*<0.0001, tp3 in **Fig. 4e**, **Supplementary Table 2**). Taken together, these results suggest that mutations in PNPO delay but do not abolish alcohol clearance.

### Mutations in the *PNPO* gene affect the biphasic response to alcohol

Alcohol induces complex behavioral responses through a variety of targets and mechanisms, including ion channels, receptors, and intracellular signaling pathways^3,6,12,40^. These mechanisms contribute to alcohol’s biphasic effect on both mood and motor controls^41–44^. Since we observed changes in sensitivity of *PNPO* mutants to alcohol toxicity, aversion and consumption, and alcohol-induced sedation (**Figs. 2-4**), we further assessed locomotor activity response of flies from each genotype to alcohol vapor exposure.

Flies were first recorded for their baseline activity for 1h and then were exposed to 95% ethanol vapor (**Fig. 5a**). Before alcohol exposure, *h^WT^* and *h^R116Q^* tended to have lower activity than other genotypes, but the difference was not significant (**Fig. 5b**, **Supplementary Table 2**). Upon alcohol exposure, flies from all genotypes showed biphasic responses such that they increased activity for the first 30min and then decreased over the subsequent 2.5h (**Fig. 5a**). We examined the peak activity and the total activity change in flies of each genotype upon alcohol exposure and found that *h^WT^* and *h^D33V^* had lower peak activity responses to alcohol than *h^R116Q^*, *sgll*^95^, and *w^1118^* (Tukey’s post hoc, *p*<0.05 for both *h^WT^* and *h^D33V^*, **Fig. 5c**, **Supplementary Table 2**). Total activity changes were similar but less pronounced (Tukey’s post hoc, *p*<0.05 for *h^D33V^*, **Fig. 5d**). These trends were consistent with genotype differences in alcohol sensitivity (**Fig. 4b-c**). In all genotypes, activity decreased to the baseline level around 2h after alcohol administration.

**Fig. 5:**
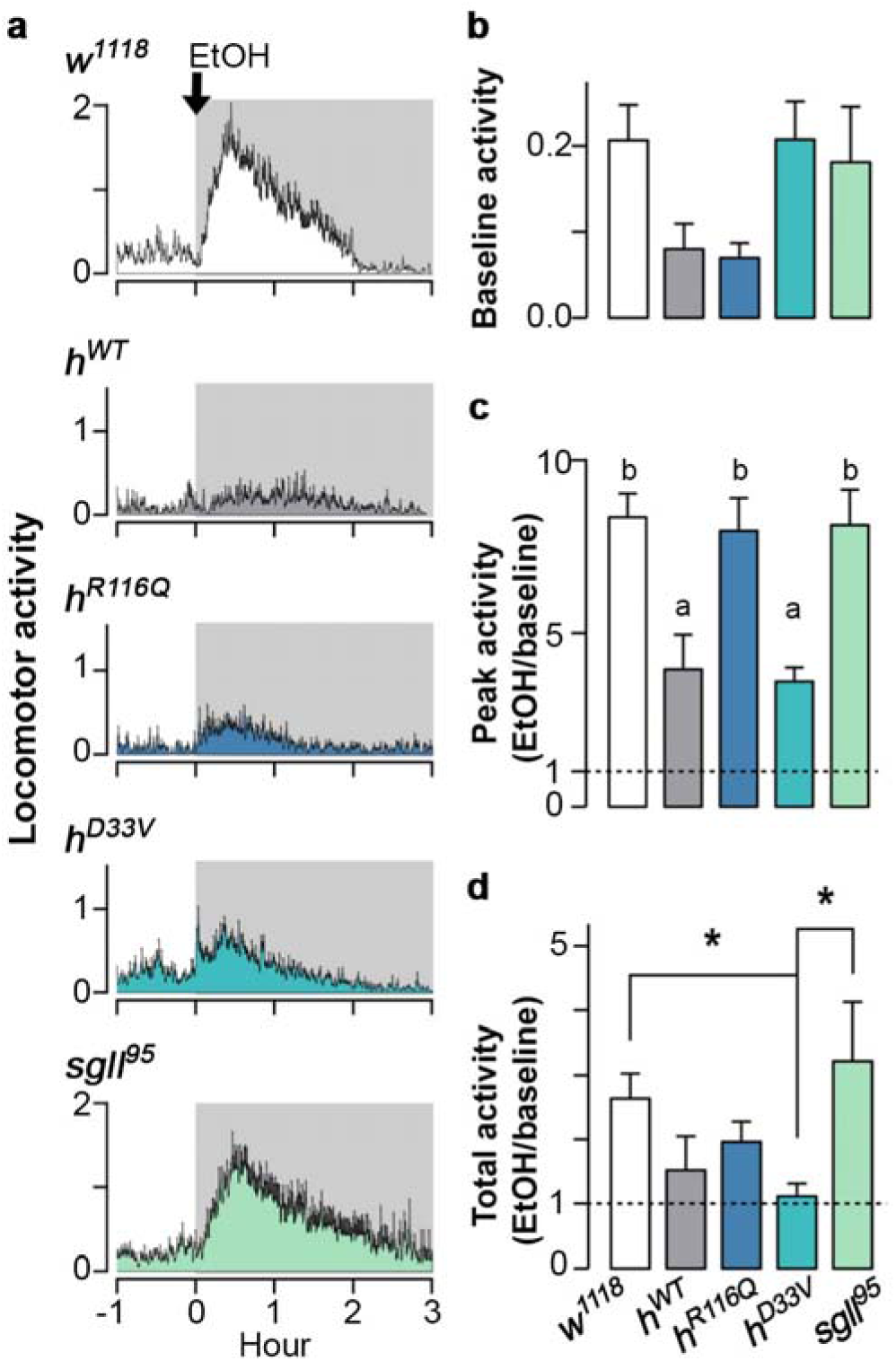
PNPO mutations affect the biphasic alcohol locomotor response. **a** Locomotor response of flies to acute alcohol exposure (shaded area). *n*=7-13 vials per genotype. **b** Baseline activity before EtOH application. **c** Fold change in peak activity after EtOH application. **d** Fold change in total activity in 3h. Data is presented as mean ± SEM. One-way ANOVA with Tukey’s post hoc. *p<0.05. In panel C, ‘a’ and ‘b’ indicate that genotypes with ‘a’ are different from genotypes with ‘b’. See Supplementary Table 2 for exact p values.

### Acute alcohol exposure and PNPO deficiency primarily converge on amino acid metabolism, altering neurotransmission

Alcohol exposure is associated with dramatic metabolic changes in humans^45^. On the other hand, PLP is a critical cofactor for more than 140 enzymes across diverse metabolic pathways^16^. To investigate the interaction between acute alcohol exposure and PNPO deficiency, we conducted metabolomic profiling of flies without or with 1h exposure to 40% ethanol vapor. We focused on the three human alleles: *h^WT^*, *h^R116Q^*, and *h^D33V^* (see Discussion). For each genotype, we triplicated samples of 60 flies and performed liquid chromatography-mass spectrometry (LC-MS, see Methods).

A total of 292 metabolites were detected, including amino acids, neurotransmitters, tricarboxylic acid cycle (TCA) intermediates, and various carnitines (**Supplementary Table 3**). To ensure data quality, we retained 290 metabolites with a coefficient of variation <30% in positive quality control samples, excluding dopamine and 2-phosphoglycerate from the final analyses. Principal component analysis (PCA) revealed clear sample separation along principal components 1 and 2 (PC1 and PC2), corresponding to alcohol treatment and genotype, respectively (**Fig. 6a**). The top 10 metabolites in PC1 included 3-hydroxybutyrate, acetyl CoA, alanine, N-acetylphenylalanine, 2-hydroyglutarate, glucose, lactate, gluconate, and N-acetylleucine, while PC2 included C7:0-DC carnitine, 5-methylcytosine, C6:0 carnitine, 3-hydroxybutyrylcarnitine, xanthosine, pterin, urate, C12:0 carnitine, C10:1 carnitine, and kynurenine (**Supplementary Table 4**).

**Fig. 6:**
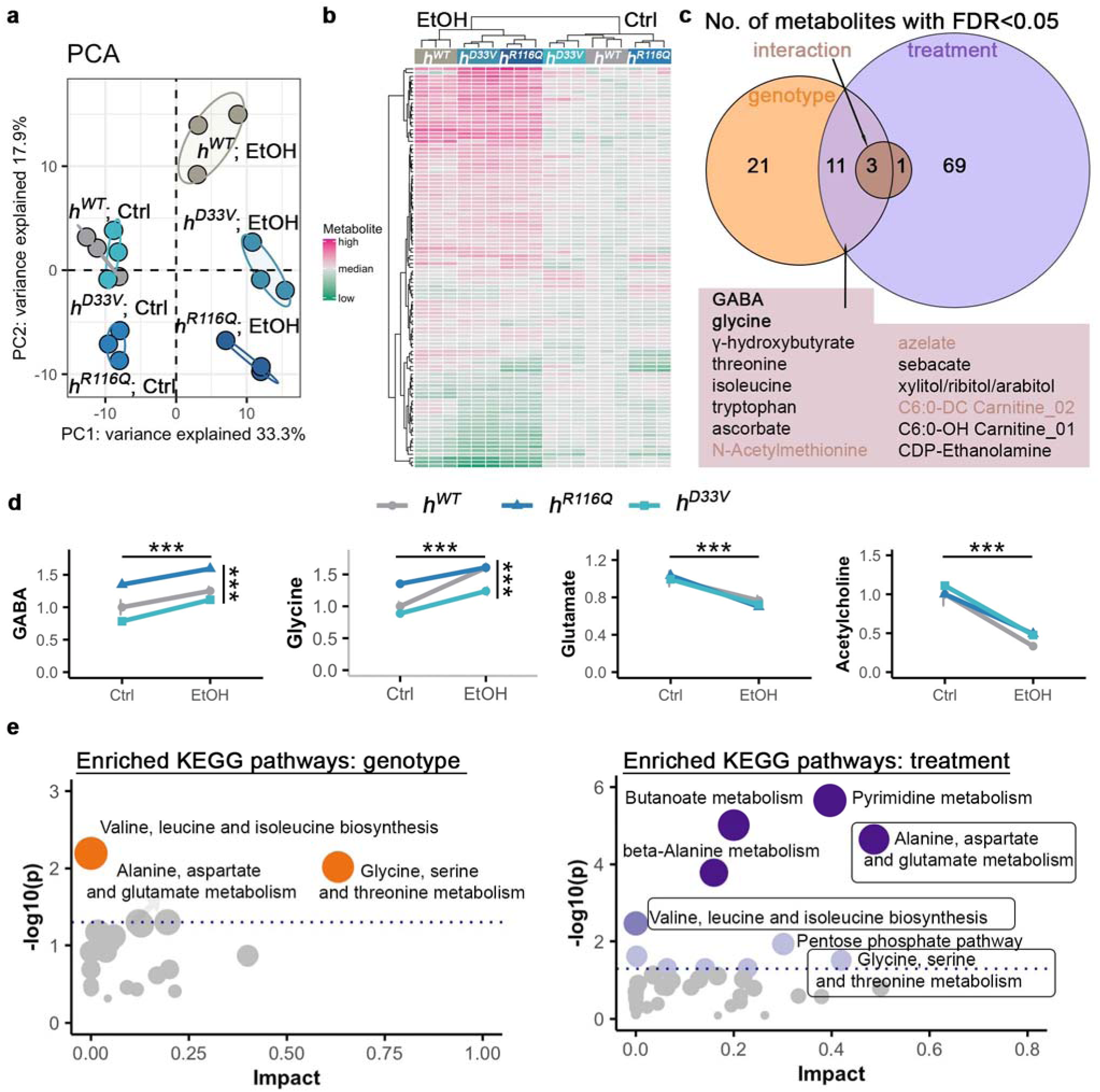
Metabolic profiling after acute alcohol exposure. **a** Principal component analyses (PCA) of metabolomics data from flies in control (Ctrl) or alcohol exposure (EtOH) conditions. **b** Heatmap of metabolites with FDR<0.05 (See Supple. Fig. 4 for metabolite information). Hierarchical clustering on both samples (columns) and metabolites (rows). **c** Venn diagram of the number of metabolites with FDR<0.05. **d** Four neurotransmitter levels from flies in Ctrl or EtOH conditions. Data is presented as mean ± SEM. *n*=3 groups per genotype per treatment condition. Two-way ANOVA with Tukey’s post hoc. The horizontal and vertical bars indicate the treatment and genotype effect, respectively. ***p<0.001. Significance for genotype comparisons within treatment groups are included in the text and exact p values are included in the Supplementary Table 2. **e** Pathway enrichment analysis for genotype and treatment. Shared pathways between genotype and treatment are in boxes. Dotted lines indicate p<0.05. See Supplementary Table 6 for complete information.

Under control conditions, *h^D33V^* clustered with *h^WT^*, whereas *h^R116Q^* samples formed a distinct group (**Fig. 6a)**, suggesting that metabolic profiles are independent of residual PNPO enzymatic activity. However, after alcohol exposure, *h^D33V^* no longer clustered with *h^WT^* but instead grouped with *h^R116Q^* (**Fig. 6b)**, indicating a potential interaction of alcohol exposure with PNPO deficiency.

To assess genotype-by-treatment interactions, we conducted two-way ANOVAs on 290 metabolites, applying a False Discovery Rate (FDR) threshold < 0.05 for statistical significance (**Supplementary Table 5**). A total of 105 metabolites were significantly affected: 35 by genotype, 84 by treatment, and 4 by genotype-treatment interaction. The four interaction metabolites were N-acetylmethionine, azelate, C6:0-DC carnitine, and fucose. Notably, 14 metabolites, including these four, overlapped between genotype and treatment effects (**Fig. 6c**), representing 13.3% of all significant metabolites. This overlap suggests that while PNPO deficiency and alcohol exposure largely affect different metabolic pathways, they converge on key metabolites, including the primary inhibitory neurotransmitter GABA.

GABA is synthesized from glutamate by GAD, which requires PLP as a co-factor^16^. Its degradation, catalyzed by GABA transaminase (GABA-T), also depends on PLP. Under control conditions, *h^D33V^* exhibited GABA levels comparable to *h^WT^*, whereas *h^R116Q^* had significant elevated GABA (Tukey’s post hoc, *p*=0.013 vs. *h^WT^*, *p*=0.0002 vs. *h^D33V^*; **Fig. 6d**, **Supplementary Table 2**). Alcohol exposure increased GABA levels across all genotypes (ANOVA, treatment main effect *F_(2,12)_*=32.56, p=9.82e-05), with *h^D33V^* showing the most significant change after multi-test correction (*p*=0.02, EtOH vs. Ctrl; **Fig. 6d**, **Supplementary Table 2**).

To identify metabolic pathways influenced by PNPO deficiency and alcohol exposure, we performed pathway enrichment analyses for non-carnitine metabolites (see Methods). Two pathways were significantly enriched for genotype and eleven for alcohol exposure (**Fig. 6e**, **Supplementary Table 6**). Notably, two genotype-associated pathways were related to amino acid metabolism, underscoring the essential role of PNPO in this process. In contrast, alcohol exposure affected more diverse pathways, including fatty acid, amino acid, and carbohydrate metabolism. Two pathways overlapped between genotype and treatment effects are: valine, leucine and isoleucine biosynthesis; glycine, serine and threonine metabolism.

Since amino acids serve as precursors for neurotransmitters, we examined inhibitory neurotransmitters (GABA, glycine) and excitatory neurotransmitters (glutamate, acetylcholine). Similar to GABA, glycine levels were elevated in *h^R116Q^* under control conditions (Tukey’s post hoc, *p*=6.47e-03 vs. *h^WT^*, *p*=6.47e-04 vs. *h^D33V^*; **Fig. 6d**, **Supplementary Table 2**). Alcohol exposure further increased glycine levels across all genotypes, with a significant genotype-by-treatment interaction (ANOVA, treatment main effect *F_(2,12)_*=82.58, *p*=9.97e-07; genotype-by-treatment interaction *F_(2,12)_*= 5.464, *p*=0.02), indicating differential genotype responses. In contrast, glutamate and acetylcholine levels were comparable among genotypes under control conditions but significantly decreased after alcohol exposure, with no significant genotype or genotype-by-treatment effects observed (ANOVA, treatment main effect, *F_(2,12)_*= 60.57, *p*=4.98e-06 and *F_(2,12)_*=125.36, *p*=1.04e-07, for glutamate and acetylcholine, respectively. **Fig. 6d**, **Supplementary Table 2)**.

Taken together, these findings demonstrate that PNPO deficiency and alcohol exposure primarily converge on amino acid metabolism, ultimately leading to increased inhibition and reduced excitation.

## Discussion

Here, we conducted behavioral and metabolomic studies to investigate the interaction between alcohol exposure and PNPO deficiency. Behaviorally, PNPO deficiency reduced aversion to high alcohol concentrations, increased alcohol consumption, and altered locomotor responses. Biochemically, PNPO deficiency and alcohol exposure primarily converged on amino acid metabolism, leading to elevated inhibitory neurotransmitters GABA and glycine. Additionally, both PNPO deficiency and alcohol exposure resulted in increased lethality in adult flies. These findings highlight a functional interaction between PNPO and alcohol exposure.

Given both PNPO deficiency and alcohol exposure can cause PLP deficiency (**Fig. 1**), this functional interaction is likely mediated through PLP. PLP is an essential cofactor for numerous enzymatic reactions, including decarboxylation, transamination, racemization, and α-elimination and replacement^16^. Decarboxylation is critical for generating neurotransmitters such as GABA, dopamine, and serotonin, while the other reactions regulate amino acid metabolism. Consistently, our metabolomic results revealed disruptions in GABA and amino acid metabolism in response to both PNPO deficiency and alcohol exposure (**Fig. 6**). Furthermore, the ability of PLP supplementation to rescue the lethality induced by PNPO deficiency and alcohol exposure supports this hypothesis (**Fig. 2**).

PNPO-deficient flies exhibited altered behavioral responses to alcohol exposure (**Figs. 3-5**). Notably, these behavioral changes did not always correlate with residual PNPO enzymatic activity, suggesting a complex relationship between PNPO mutations and alcohol-related behaviors. This complexity may arise because each behavioral assay captures a distinct aspect of alcohol response, governed by specific signaling pathways or neural mechanisms^22,39^. Additionally, mutation-specific properties may contribute to these differences. For example, while biochemical studies indicate that the R116Q mutation is milder than D33V^17^, we previously found that R116Q mislocalized PNPO from the soma to terminal structures in the adult fly brain^24^. This mislocalization likely alters the subcellular availability of PLP, thereby affecting PLP-dependent enzymes.

In GABA synthesis, altered PNPO subcellular localization could lead to changes in synaptic GABA levels, as the two forms of GAD, GAD67 and GAD65, have distinct subcellular localizations. GAD67 is primarily localized in the soma and is constitutively bound to PLP, whereas GAD65 is localized at synaptic terminals and its activity is tightly regulated by PLP availability^46,47^. Thus, increased PNPO localization at synaptic terminals in *h^R116Q^* flies may lead to enhanced GABA synthesis at synaptic sites. Supporting this, we observed significantly elevated GABA levels in *h^R116Q^* flies (**Fig. 6D**), although future studies are needed to determine whether this increase is specific to synaptic terminals. The elevated GABA levels in *h^R116Q^* compared to *h^WT^* and *h^D33V^* may underlie the significantly increased peak locomotor activity of *h^R116Q^* flies in response to alcohol exposure (**Figs. 5-6**). These findings underscore the importance of *in vivo* studies to fully elucidating the functional consequences of disease-associated mutations beyond their effects on residual enzymatic activity.

*Drosophila sgll* has two splicing isoforms RA and RB, with RB being the predominant and widely expressed form^24^. In our humanized PNPO model (*h^WT^*), the RB isoform was replaced by human *PNPO* cDNA. The fact that behavioral responses of *h^WT^* were not always consistent with those of *w^1118^* (**Fig. 4C, 5C**) suggests that the RA isoform may have functional significance. Alternatively, differences between *h^WT^* and *w^1118^* could stem from sequence divergence between human and fly PNPO, which share only 45% identity.

Intriguingly, we observed a significant genotype-by-treatment interaction for glycine (**Fig. 6**), an amino acid that is involved in one-carbon metabolism and functioning as an inhibitory neurotransmitter. One-carbon metabolism encompasses biochemical pathways that transfer and utilize one-carbon (methyl) groups for essential cellular processes, including nucleotide synthesis and epigenetic regulation^48^. Consistently, *PNPO* gene knockdown in flies induces chromosome aberrations^49^. Future studies could explore whether PNPO mutant flies exhibit deficits in epigenetic regulation.

In humans, excessive pyridoxine intake induces peripheral neuropathy, characterized by symmetric, progressive impairments in touch, pain, temperature, vibration, and proprioception in the extremities^27,28^. Despite well-documented symptoms, the specific VB6 vitamer responsible remains unclear due to the interconversion among VB6 vitamers^28^. Our study provides compelling evidence that high PLP levels are toxic (**Fig. 2**). As a highly reactive aldehyde, PLP may exert its toxicity through multiple mechanisms, including aldehyde-mediated damage^28^.

AUD affects 10-15% of the global population, imposing substantial medical, social, and economic burdens^50,51^. Current AUD treatments primarily target ethanol metabolism and neurotransmitter systems^52^. Our findings suggest that PNPO deficiency may contribute to AUD and alcohol toxicity and that VB6 supplementation at appropriate doses may represent a potential therapeutic strategy for AUD, particularly in individuals with genetic deficiency in VB6 metabolism.

## Materials and Methods

### *Drosophila* strains and husbandry

All fly strains used in this study were previously described^24,25^. Flies were generated on the standard cornmeal-yeast-molasses or Frankel & Brosseau’s media. Flies used in all experiments were raised and tested at ∼23°C under a 12:12-h light:dark cycle. Male flies, 1 to 3 d old, were picked and used in the experiments corresponding to Figures 1, 2, 3, 4, and 6. Male flies, 1 to 7 d old, were picked and used in the experiment corresponding to Figure 5.

### VB6 vitamer measurement

For vitamin B6 species analysis, 60 snap-frozen flies per genotype were collected either without alcohol exposure or after 1h exposure to 40% EtOH. VB6 species were extracted using ice-cold 10% trichloroacetic acid (8µL solvent per mg of fly tissue, LC-MS grade water), homogenized (Omni International, TH115-PCR5H, stainless steel probe), vortexed, sonicated for 3min in an ice-cold water bath, and vortexed again for 5min at 2,000rpm and 4°C. After incubation on ice for 20min, samples were centrifuged at 20,000g for 25min at 4°C, and 75µL of supernatant was transferred to an LC-MS vial. A pooled quality control sample was created by combining ∼20uL from each sample and injected every 9th run to assess reproducibility. Measurements were performed using LS-MS at the Metabolomics Core Facility at the University of Chicago.

### Lifespan and survival assay

Groups of 15-20 flies per genotype were housed in polypropylene vials containing 4% sucrose in 1% agar with either 16% EtOH (Sigma-Adrich #EX0290) or an EtOH-free control. Additional cohorts were supplemented with 0, 4, 40, or 400 µg/mL PLP (Sigma-Aldrich #P9255). Survival was recorded daily for 21 days.

### Alcohol preference

Alcohol preference was assayed using proboscis extension response (PER) and a binary food choice assay^34,35^. For PER: 18-40 flies per genotype were starved for 16 hours, then individually placed head-first into a pipette tip, leaving the head and forelimbs exposed. A 6mm strip of Kimwipe paper was twisted into a thread and pulled apart into cone-shaped wicks. These wicks were dipped in either 16% EtOH + 4% sucrose or an isocaloric maltodextrin control solution with 4% sucrose and then brought to the flies’ labellum to assay proboscis extension. To rule out incorrect stimulation, each *Drosophila* was presented with the EtOH- and maltodextrin-soaked wick up to 3 times. The number of responsive flies was recorded, and the assay was repeated 3 times. For binary food choice assay, flies were given a choice between 16% EtOH or isocaloric maltodextrin (Sigma-Aldrich #419672) mixed with 4% sucrose and 1% food dye (FD&C Blue No. 1 or Red No. 40; Spectrum) in 3% agar. Food was placed in alternating wells in a 96-well microplate (Corning, Falcon #353072). After 16h of starvation, 25-30 flies per genotype were placed in the plate within a dark chamber for 60min. Flies consuming EtOH or control food were identified by abdomen color. Preference index was calculated as: (Number of flies eating EtOH food)-(Number of flies eating control food)/(Number of flies eating EtOH food)+(Number of flies eating control food)+(Number of flies eating both). The assay was repeated 8 times for EtOH paired with blue dye and 9 times for EtOH paired with red dye.

### Alcohol sensitivity and tolerance

Sensitivity and tolerance were assessed as described previously^39^. Briefly, 15 flies were transferred to food vials 24 hours before the assay and then placed in empty vials sealed with a cotton swab soaked in 3mL of 40% EtOH in H_2_O with 1% FD&C Blue No. 1 dye. Sedated flies (falling to the vial bottom) were counted every 6min for 1h. Flies were then transferred into fresh vials for recovery (16h), and non-recovered flies were removed. Flies were subsequently re-exposed to 40% EtOH, and sedation was recorded every 6min for 1.5h. The time to sedation of 50% of flies (ST50) for both exposures was calculated with data fitting a log-logistic model with R functions drm and ED^53^. Tolerance was calculated as the ST50 ratio between the second and the first exposures.

### Tissue alcohol content

EtOH metabolism was assessed using a modified protocol^54,55^. Briefly, 15 flies exposed to 1h of 40% EtOH were homogenized in 75µL of 100mM Tris-HCl (Sigma-Aldrich #10812846001), pH 7.5, using a Kontes Microtube Pellet Pestle (DWK Life Sciences #749540-0000). Homogenates were mixed with 40µL of 3.4% perchloric acid (Sigma-Aldrich #244252), centrifuged at 2,000rpm for 6min at 4°C and 7µL of supernatants was mixed with 343µL of reaction buffer containing 0.5M Tris-HCl pH 8.8, 2.75µg/mL alcohol dehydrogenase (Sigma-Aldrich #A7011), and 0.5mg/mL β-nicotinamide adenine dinucleotide (Sigma-Aldrich #N7132). EtOH standards (200mM, 100mM, 50mM, 25mM, 12.5mM, 6.25mM, 3.13mM, 1.56mM, 0.78mM) were prepared similarly. Samples were incubated at room temperature for 40min, and absorbance was measured at 355nm using a VICTOR Nivo multimode spectrophotometer (PerkinElmer). EtOH concentration was calculated using the standard curve. The assay was repeated 4 times.

### Locomotor response to alcohol

Fly activity was monitored using a Drosophila activity monitor (LAM25H-3, TriKinetics Inc, Waltham, MA), with 7-10 flies per vial at 23°C under a 12:12-h light:dark cycle (Light on = 09:00 – 21:00). After overnight acclimation, 500µL of 95% EtOH in H_2_O was added to the cotton plug between 15:00 and 15:30. Activity was recorded every minute, and data were pooled from 7-13 vials to generate histograms. Peak locomotor counts were determined using a 10min running average. Baseline activity is averaged counts before EtOH treatment, calculated as counts per min per fly. Peak activity is the ratio between peak counts during EtOH and baseline activity, both of which were calculated as counts per min per fly. Total activity is the ratio between the average counts in the presence of EtOH and the average baseline activity, both of which were calculated as counts per min per fly.

### Metabolomic sample preparation, measurement, and data analysis

Groups of 60 flies per condition (with or without 1h of 40% EtOH exposure) were snap frozen. Samples were extracted with 4:4:2 acetonitrile/methanol/water (20µL solvent per mg of fly tissue, LC-MS grade), homogenized, vortexed, and subjected to two freeze-thaw cycles in liquid nitrogen. Samples were then sonicated for 3min in an ice-cold water bath, vortexed for 5min at 2,000rpm at 4°C, incubated on ice for 20min, and centrifuged at 20,000g for 20min at 4°C. Supernatant (300µL) from each sample were dried using a Genevac EZ-2.4 elite evaporator and stored at −80°C until analysis. A pooled quality control sample was injected throughout the run to ensure reproducibility. Metabolomic profiling was performed at the University of Chicago Metabolomics Core Facility. A total of 292 metabolites were detected. Final analysis included 290 metabolites that had a coefficient of variation <30% from positive quality controls. Data analysis was performed in R^56^. Enrichment analysis was done in MetaboAnalyst^57^. Fatty acid-associated carnitines were not included in the enrichment analysis due to their unavailability in the metabolite database. Pathway annotation was based on KEGG Drosophila melanogaster metabolite pathways.

### Statistical Analysis

Statistical analyses were performed in R (version 4.1.2). Details on statistical analyses, including sample sizes, tests performed, and multiple tests correct, if necessary, are provided in the text/figures/figure legends. Exact sample sizes and p values are provided in Supplementary Tables 1 and 2.

## Acknowledgments

We thank Amy Lasek for sharing her alcohol tissue content assay. We thank Abigail Hawken, Bryan Joos, and Kellen M Ochs for collecting the locomotor activity data in Figure 5. This work was supported by NIH Grants T32DA043469 (to W.C. and B.W.) and R01NS111122 (to X.Z.).

---

**Supple. Fig. 1:**
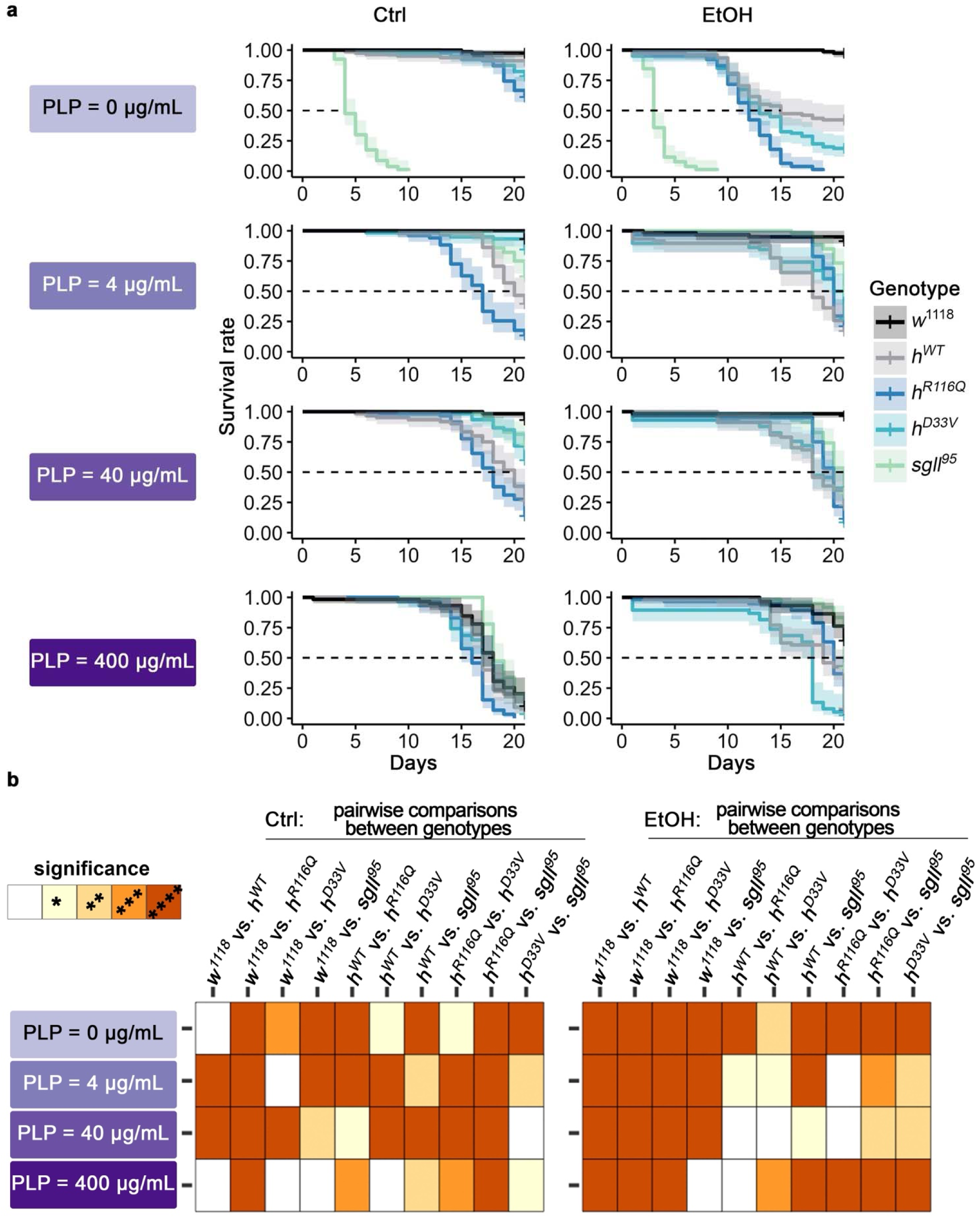
Effects of chronic alcohol exposure and PLP supplementation on survival of flies from each genotype. **a** Survival of flies from each genotype on control diet (Ctrl) or 16% alcohol (EtOH) for 21 days in combination with PLP supplementation. *n*=38-80 flies per genotype per diet treatment. Log-rank test with Benjamini-Hochberg corrections. **b** Pairwise log-rank test with Benjamini-Hochberg corrections. *p<0.05, **p<0.01, ***p<0.001, ****p<0.0001. See Supplementary Tables 1-2 for exact n in each condition and exact p values for comparisons.

**Supple. Fig. 2:**
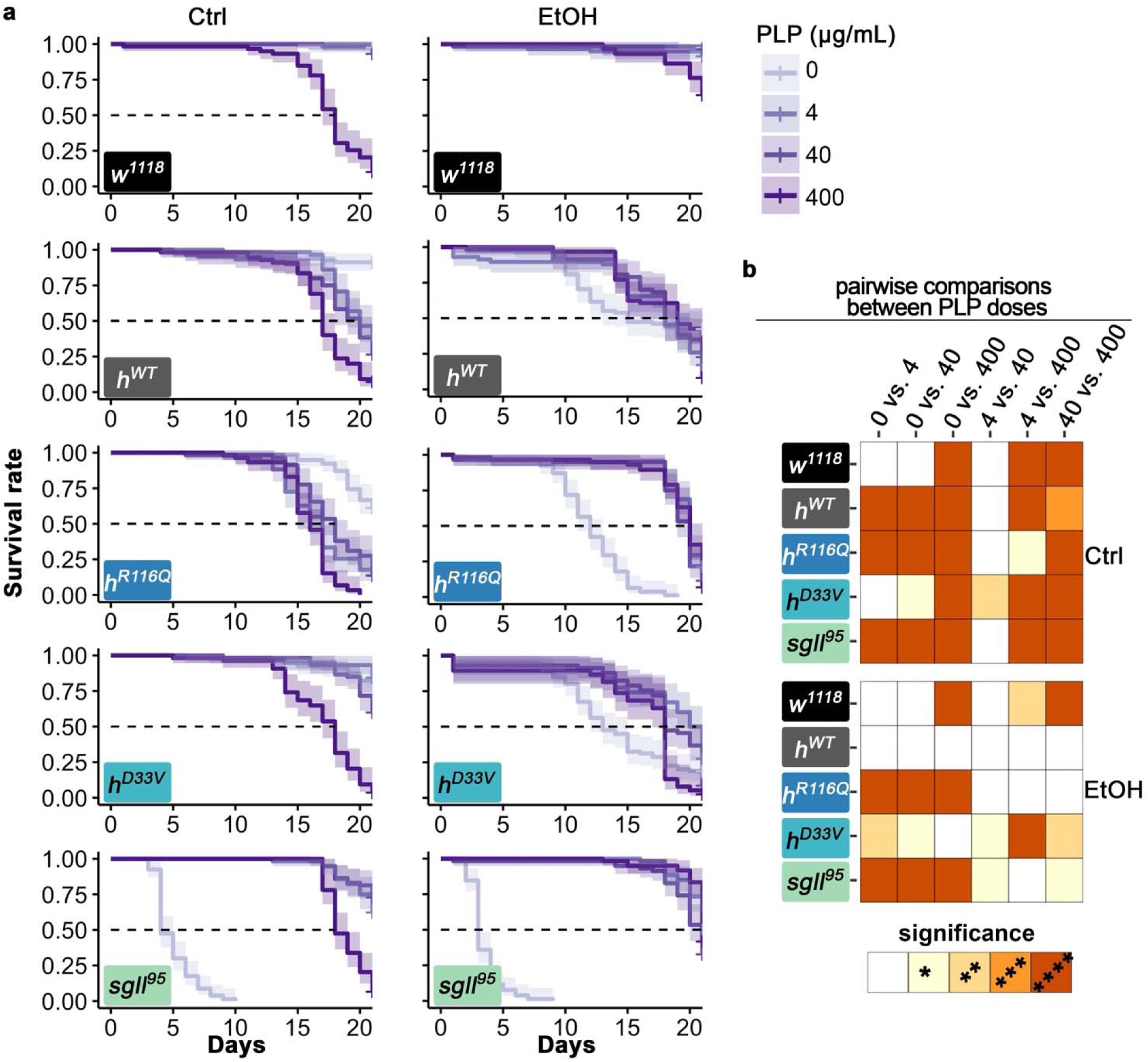
Effects of chronic alcohol exposure and PLP supplementation on survival of flies from each genotype. **a** Survival of flies from each genotype on control diet (Ctrl) or 16% alcohol (EtOH) for 21 days in combination with PLP supplementation. *n*=38-80 flies per genotype per diet treatment. **b** Pairwise log-rank test with Benjamini-Hochberg corrections. *p<0.05, **p<0.01, ***p<0.001, ****p<0.0001. See Supplementary Tables 1-2 for exact n in each condition and exact p values for comparisons.

**Supple. Fig. 3:**
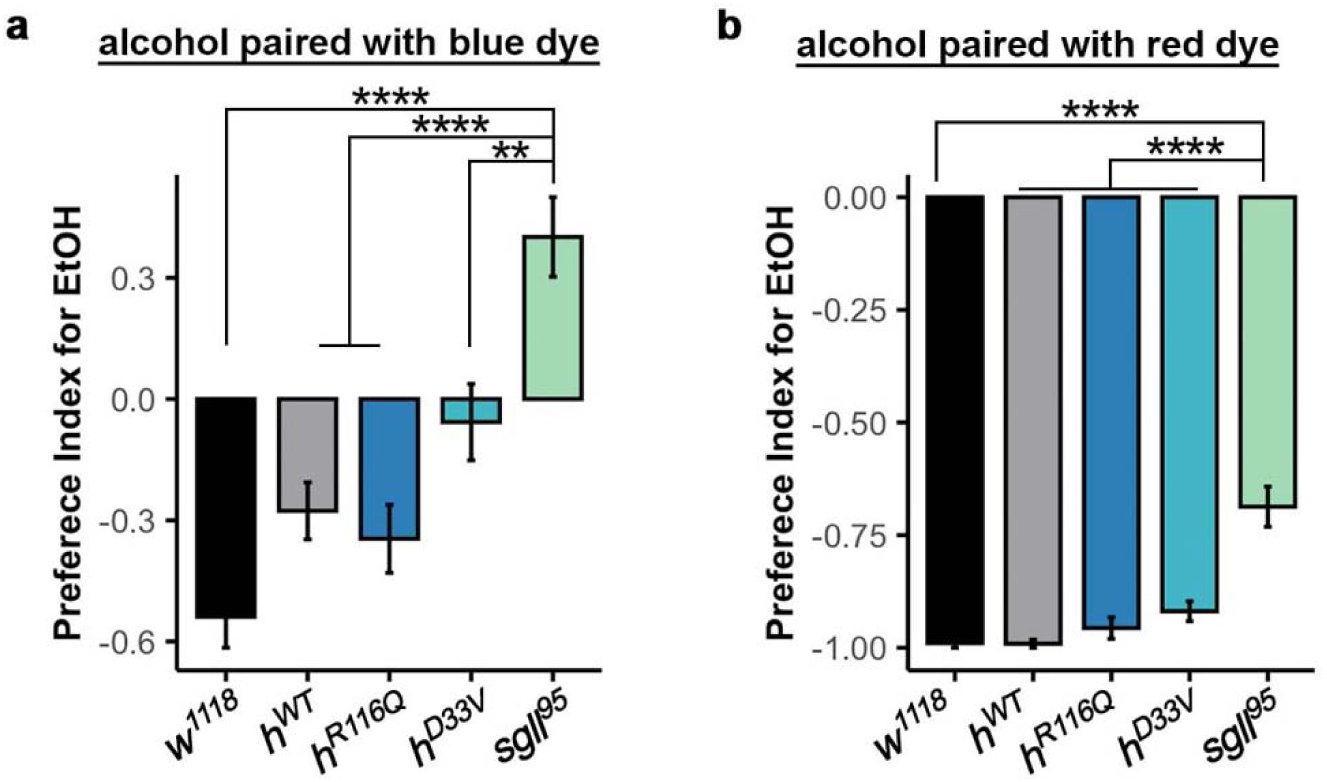
Preference Index for EtOH over isocaloric maltodextrin. *n*=8 and 9 assays per genotype for alcohol paired with blue dye and paired with red dye, respectively. Data is presented as mean ± SEM. One-way ANOVA with Tukey’s post hoc. n.s.:p>0.05, *p<0.05, **p<0.01, ***p<0.001, ****p<0.0001. See Supplementary Table 2 for exact p values.

**Supple. Fig. 4.**
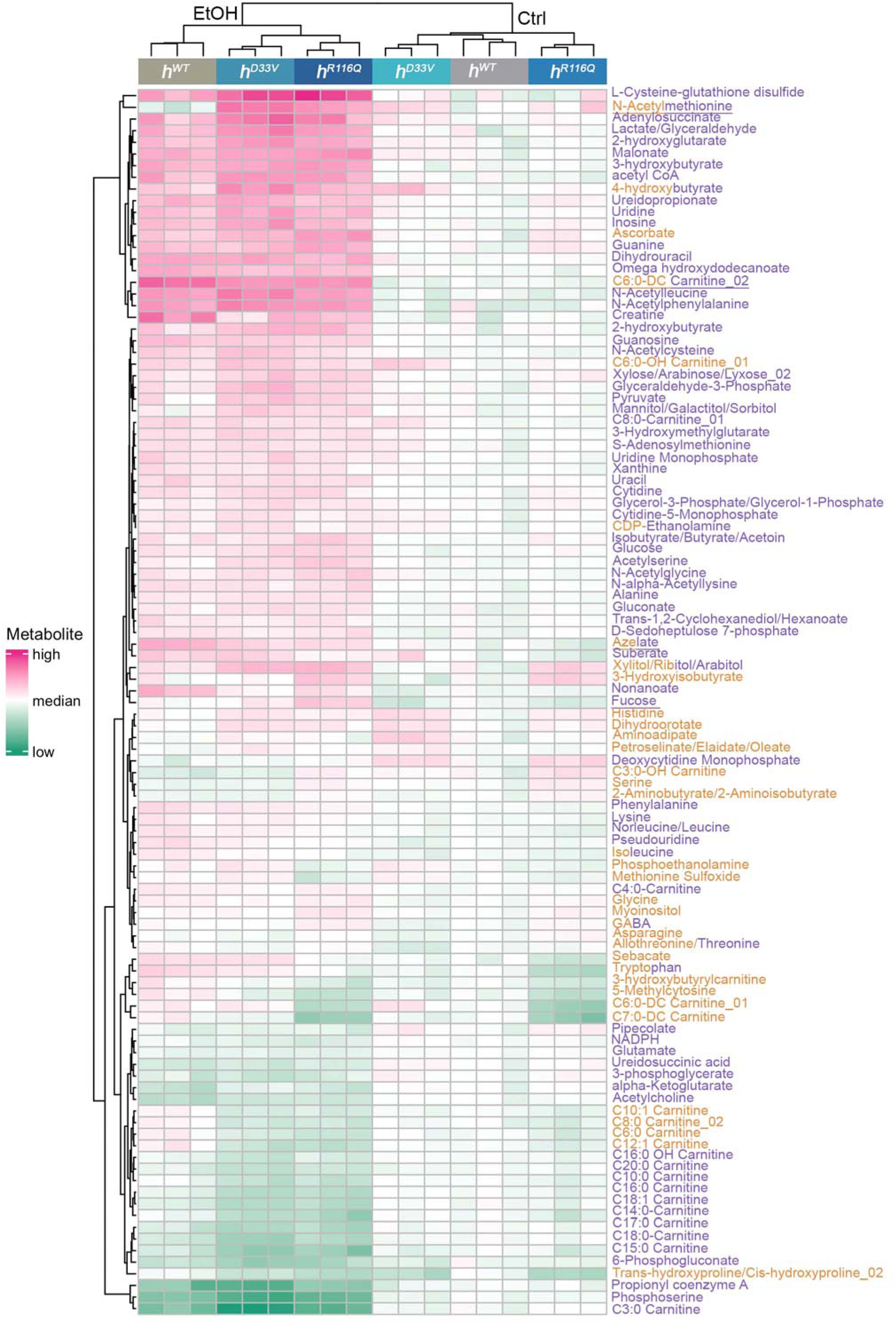
Hierarchical clustering of metabolites. Only metabolites with FDR<0.05 are included. Hierarchical clustering on both samples (columns) and metabolites (rows). Metabolites are color-coded. Orange and purple indicate significant metabolites for genotype and treatment, respectively. Metabolites with dual colors are significant for both genotype and treatment. Metabolites with underlines are significant for genotype-by-treatment interaction.

